# Lack of neutralizing antibodies against the current circulating influenza viruses during the Omicron outbreak in Hong Kong

**DOI:** 10.1101/2023.02.07.527570

**Authors:** Weiwen Liang, Huibin Lv, Chunke Chen, Yuanxin Sun, David S Hui, Chris Ka Pun Mok

**Affiliations:** HKU-Pasteur Research Pole, School of Public Health, Li Ka Shing Faculty of Medicine, The University of Hong Kong, Hong Kong SAR, China; Department of Biochemistry, University of Illinois at Urbana-Champaign, Urbana, IL 61801, USA; The Jockey Club School of Public Health and Primary Care, The Chinese University of Hong Kong, Hong Kong SAR, PR China; Li Ka Shing Institute of Health Sciences, Faculty of Medicine, The Chinese University of Hong Kong, Hong Kong SAR, PR China; Department of Medicine and Therapeutics, The Chinese University of Hong Kong, Hong Kong SAR, PR China; Stanley Ho Centre for Emerging Infectious Diseases, Faculty of Medicine, The Chinese University of Hong Kong, Hong Kong SAR, China

## Abstract

We report the seroprevalence to the circulating influenza A H1N1 and H3N2 viruses from plasma samples collected from 479 adults between 2021 and 2022. Our results show that there is lack of neutralizing antibodies to these viruses and highlight the importance of promoting influenza vaccination during the emerging of COVID-19.

## Background

As the Omicron variants of SARS-CoV-2 are now persistent worldwide, “Living with COVID-19” has been adopted by many countries especially those can achieve high coverage rate of COVID-19 vaccination. Preventive measures such as wearing mask and keeping social distancing are no longer compulsory. The prevalence of influenza A viruses remains low during the COVID-19 outbreak. However, there is a concern that lifting of the stringent preventive policies may result in the widespread of seasonal influenza viruses. Importantly, recent studies showed that co-infection of SARS-CoV-2 and influenza virus was associated to more severe symptoms in humans (Adams et al., 2022; Pawlowski et al., 2022). Nevertheless, there is no serological study so far to determine the levels of neutralizing antibodies against recent circulating influenza strains after the COVID-19 outbreak emerged.

## The Study

From 23^rd^ July 2020 to 31^st^ December 2022, only 344 influenza infected cases had been recorded through the weekly influenza surveillance in Hong Kong (https://www.chp.gov.hk/en/resources/29/304.html). In this study, we tested the plasma samples collected from adults in Hong Kong and determined the levels of neutralizing antibodies against the H1N1 virus namely A/Wisconsin/588/2019 (Wis19) and H3N2 virus A/Darwin/6/2021 (Dar21), which are the compositions of seasonal influenza vaccine for the 2022-2023 season. The hemagglutinin sequences of both viruses are phylogenetically closed to the circulating strains which have been recently isolated worldwide including the isolates from Hong Kong **(Figure 1)**. Paired plasma samples from 479 adults were collected at two time points (2021 and 2022) between 8^th^ March 2021 and 21^st^ October 2022. Samples at the 2^nd^ time point were collected in 2022 except one was obtained on 20^th^ Dec 2021. Among all participants, 283 adults reported that they had not received influenza vaccine before their first sampling and 196 adults reported that they received the vaccine annually. The mean intervals between the 1^st^ and 2^nd^ sampling time point were 420.15 (No vaccination) and 442.84 days (Vaccination annually), respectively. The demographic information and the microneutralization results are shown in **Table 1** and **Supplementary Figure 1**. We defined the samples as negative if the microneutralization titers were lower than 1:20 which was the lower detection limit in our experiment. Among those who had not taken influenza vaccine, the positive rates of neutralizing antibodies against H1N1/Wis19 and H3N2/Dar21 in 2022 were only 5.30% (15/283) and 0.35% (1/283), respectively. Although 28.06% (55/196) of participants who reported receiving influenza vaccine annually were found as seropositive to H1N1/Wis19, only 1 sample in this group had neutralizing titer against H3N2/Dar21 equal to or higher than 1:20. As controls, the levels of neutralizing antibodies against A/Michigan/45/2015 (Mich15) and A/Singapore/INFIMH-16-0019/2016 (Sing16) which were the compositions of flu vaccine for the 2018-2019 season were also measured. As expected, the neutralizing antibodies against both viruses in annual vaccination group were higher than those who reported without receiving influenza vaccination **(Table 1)**. Surprisingly, while the overall protection against H3N2/Dar21 was negligible in both groups, there was a significantly higher positive rate of neutralizing titer against H1N1/Wis19 in those who received vaccine previously comparing to the no vaccine group (Table 1). Further analysis showed that the levels of neutralizing titers against the H1N1/Mich15 and H1N1/Wis19 were statistically correlated, suggesting that the antibodies against H1N1/Mich15 might partially cross-react to recent circulating H1N1 strains **(Figure 2)**.

**Figure 1.**
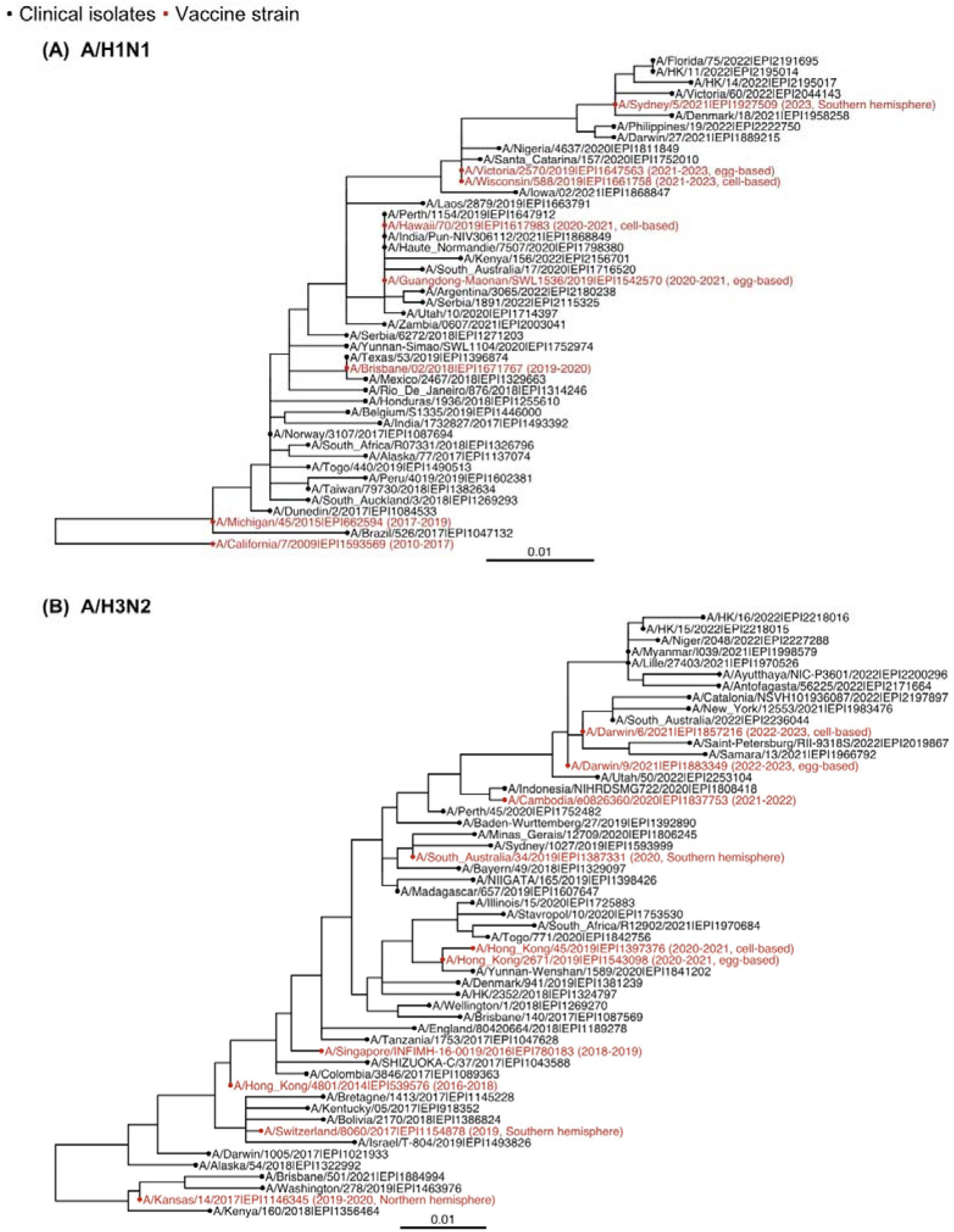
Phylogenetic trees of seasonal influenza/A H1N1 and H3N2 viruses. Evolution of seasonal influenza A viruses is shown by two rooted phylogenetic trees of **(A)** H1N1 and **(B)** H3N2 viruses using representative strains isolated between January 2017 and 2022. Vaccine strains are indicated in red while circulating wild type strains are indicated in black.

**Figure 2.**
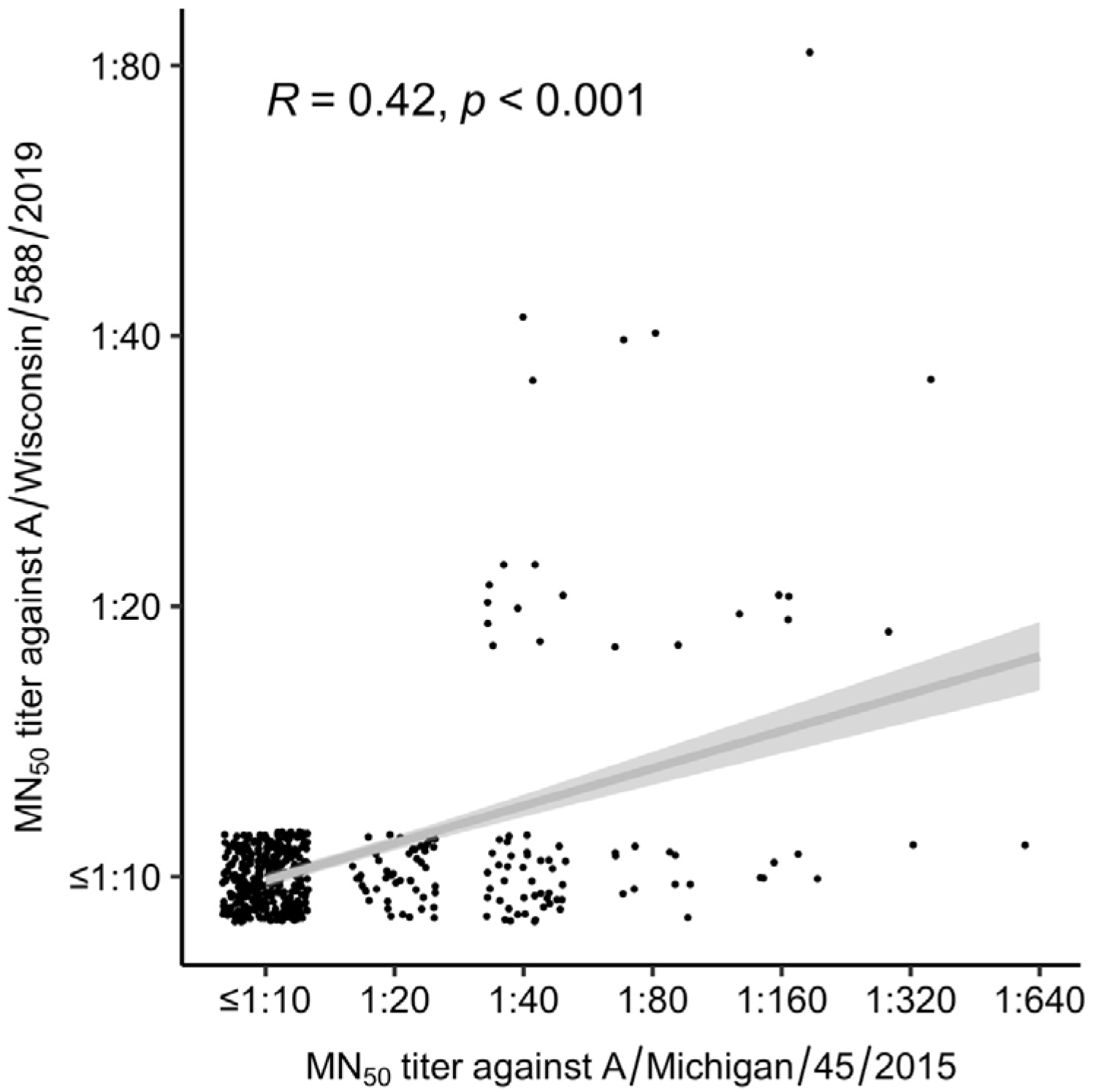
Correlation between the neutralizing titers against H1N1 viruses. The correlation of the results from microneutralization (MN) titers that inhibits at least 50% infection of H1N1/Mich15 and H1N1/Wis19 were determined. The Spearman’s rank coefficient (*R* value) and *p* value were used to indicate the correlation between MN_50_ results.

## Conclusion

Our results suggest that adults in the Hong Kong community generally lack sufficient neutralizing antibodies against circulating influenza strains. Among all participants (n=479), only 2 (0.42%) and 70 (14.61%) had neutralizing antibodies against H3N2/Dar21 and H1N1/Wis19 viruses, respectively. Previous vaccination may contribute to the protection against the new H1N1 but not the new H3N2 virus. The neutralizing antibody against the H1N1/Wis19 is likely contributed by the cross-reactive immunity to the previous H1N1 strain. It is expected that the compositions of influenza A viruses which are used for 2022-2023 vaccine match the recent circulating strains. Thus, influenza vaccination should be actively promoted, especially to the high risk groups like elderly or immunocompromised patients.

## Acknowledgments

This research was supported by grants from the Health and Medical Research Fund Commissioned Research on the Novel Coronavirus Disease (COVID-19), Hong Kong SAR (COVID1903003), Emergency Key Program of Guangzhou Laboratory (Grant No. EKPG22-30-6), RGC’s Collaborative Research Fund (C6036-21GF), UGC Research Matching Grant (8601629) and visiting scientist scheme from Lee Kong Chian School of Medicine, Nanyang Technological University, Singapore (CKPM). We also thank the S.H. Ho Foundation for providing funding support.

## Materials and Methods

### Construction of phylogenetic tree

Full-length protein sequences of hemagglutinin (HA) from pandemic2009 (pdm09) H1N1 and H3N2 viruses isolated between 2017 and 2022 were downloaded from the Global Initiative for Sharing Avian Influenza Data (GISAID; http://gisaid.org) (Shu & McCauley, 2017). Besides, sequences of vaccine strains used from 2017 to date were also included. Sequences without any ambiguous amino acid were aligned using MAFFT v7.453 with default setting (Katoh et al., 2002). A total number of 41 and 53 strains including the vaccine viruses were selected as representative strains by CD-HIT (Li & Godzik, 2006) and used for building phylogenetic trees by IQ-TREE2 with default parameters (Minh et al., 2020). Phylogenetic trees were finally visualized using ggtree for R (Yu, 2020).

### Cell culture

Humanized Madin-Darby canine kidney (hMDCK) cells (Takada et al., 2019) and human embryonic kidney (HEK) 293T cells were used in this study and maintained in minimal essential medium (MEM) supplemented with 10% fetal bovine serum, 25 mM HEPES, and 100 U mL^-1^ penicillin-streptomycin (PS).

### Virus rescue using reverse genetics

All influenza A viruses used in this study were generated based on the A/PR/8/34 (PR8) influenza eight-plasmid reverse genetics system using the HA and NA from strains of interest and six internal genes from PR8 as backbone (Hoffmann et al., 2000; Neumann et al., 1999). Both HA and neuraminidase (NA) genes of the strains of interest were synthesized by Sangon Biotech, and cloned into the pHW2000 vector as previously described (Liang et al., 2022). Briefly, co-culture of HEK 293T and hMDCK cells at a 6:1 ratio were prepared one day before the transfection. For rescuing one 6:2 recombinant, 1μg each of the eight plasmids were mixed well with 16 μL of TransIT-LT1 (Mirus Bio) and transfected into the hMDCK-HEK293 cocultured cells at 70% confluence. Medium was replaced with 1 mL of MEM at 6 h post-transfection, and another 1 mL of MEM supplemented with 1 μg mL-1 tosylphenylalanyl chloromethyI ketone (TPCK)-trypsin (Sigma) was added at 24 h post-transfection. Supernatant was harvested at 72 h post-transfection and further inoculated into hMDCK cells at 90% confluence. Supernatant of the hMDCK cells was harvested when more than 70% of cells had cytopathic effect (CPE) and processed to viral extraction using QIAgen viral extraction kit (Thermofisher) and sanger sequencing on HA and NA to confirm the virus identity. The TCID_50_ titer was determined by titration as previously described in hMDCK cells (Chan et al., 2013; Chan et al., 2010).

### Microneutralization assay

Microneutralization (MN) assay was performed according to the manual for serological diagnosis from the World Health Organization (Organization). Briefly, approximately 2×10^4^/well of hMDCK cells in each well were prepared one day before MN assay in 96-well cell culture plates and cultured until a 100% confluent monolayer was obtained. Cells were washed once with phosphate-buffered saline (PBS) and resupplied with MEM containing 25 mM HEPES, and 100 U mL^-1^ PS. All plasma for MN assay were heat-inactivated at 56 °C for 30 min before the experiment. A two-fold serial dilution was performed on heated plasma to make series dilutions from 1:20 to 1:2560, which were subsequently mixed with 100 TCID_50_ of viruses in equivalent volume and incubated at 37 °C for 1 h. After 1-hour incubation, the mix was further inoculated into the cells and incubated at 37 °C for another 1 h. Cell supernatants were then discarded and replaced by MEM with 25 mM HEPES, 100 U mL^-1^ PS and 1 μg mL-1 TPCK-trypsin. Plates were inoculated at 37 °C for 72 h. By 72-h incubation, plasma dilution that inhibited the virus growth and CPE was recorded as the MN_50_ titer.

## Figure Legends

**Supplementary Figure 1.**
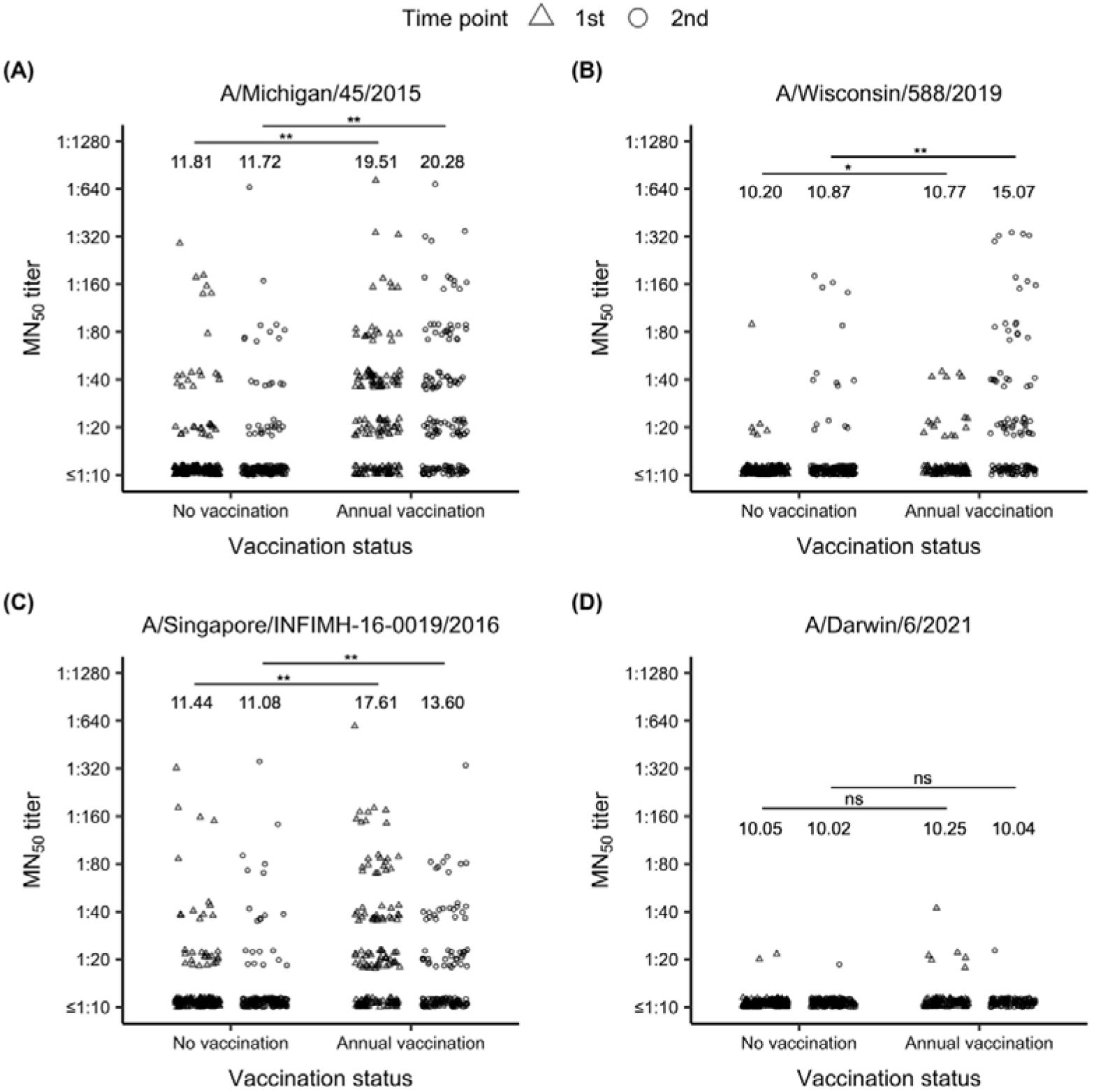
Neutralization titers against the influenza viruses. The microneutralization (MN) titers that inhibits at least 50% infection of two H1N1 influenza A viruses **(A-B)** and two H3N2 influenza A viruses **(C-D)** are shown. Paired plasma samples were collected at two time points from each adult (n= 479). Samples were divided into two groups who reported with (n= 196) or without (n=283) receiving influenza vaccination annually. MN_50_ titer ≥1:20 was defined as positive. Chi-square test was used to examine the statistical difference of positive rate between participants from the two groups. ** *p*□<□0.001; * *p*<□0.05; ns non-significant.

## Table 1

**Supplementary Table 1.** Demographic information of the participants and the neutralization results.

|  | Status of receiving influenza vaccine |  | *p* value |
| --- | --- | --- | --- |
|  | Annually (n=196) | No vaccination (n=283) |  |
| Age (Mean±SD) (Min, Max) | 50.85±11.66 (19, 79) | 47.92±12.38 (18, 77) | 0.061 |
| Gender (male, %) | 34.18% | 39.22% | 0.262 |
| Date of collection (1<sup>st</sup> time point) | 8 Mar 2021 - 27 Aug 2021 | 9 Mar 2021 - 17 Sep 2021 | - |
| Date of collection (2<sup>nd</sup> time point) | 26 Jan 2022 - 7 Oct 2022 | 20 Dec 2021 - 21 Oct 2022 | - |
| Interval between 1<sup>st</sup> and 2<sup>nd</sup> time point (Mean±SD) | 442.84 ± 57.59 days | 420.15 ± 57.87 days | <0.001 |
| Reference virus | Number of samples with MN<sub>50</sub> ≥1:20 (%) |  | *p* value |
| Mich15 (1<sup>st</sup> time point) | 95 (48.47%) | 35 (12.37%) | <0.001 |
| Mich15 (2<sup>nd</sup> time point) | 94 (47.96%) | 34 (12.01%) | <0.001 |
| Wis19 (1<sup>st</sup> time point) | 16 (8.16%) | 6 (2.12%) | 0.002 |
| Wis19 (2<sup>nd</sup> time point) | 55 (28.06%) | 15 (5.30%) | <0.001 |
| Sing16 (1<sup>st</sup> time point) | 81 (41.33%) | 31 (10.95%) | <0.001 |
| Sing16 (2<sup>nd</sup> time point) | 51 (26.02%) | 21 (7.42%) | <0.001 |
| Dar21 (1<sup>st</sup> time point) | 6 (3.06%) | 2 (0.71%) | 0.068 |
| Dar21 (2<sup>nd</sup> time point) | 1 (0.51%) | 1 (0.35%) | 1 |

Mich15: A/Michigan/45/2015; Wis19: A/Wisconsin/588/2019; Sing16: A/Singapore/INFIMH-16-0019/2016; Dar21: A/Darwin/6/2021
Differences of gender and seropositive with MN<sub>50</sub> ≥ 1:20 between the two groups were determined by Chi-square test while age and interval of sampling were compared by Mann-Whitney U test.

## References

1. Shu Y, McCauley J. GISAID: global initiative on sharing all influenza data - from vision to reality. Euro Surveill. 2017 Mar 30;22(13).

2. Katoh K, Misawa K, Kuma K, Miyata T. MAFFT: a novel method for rapid multiple sequence alignment based on fast Fourier transform. Nucleic Acids Res. 2002 Jul 15;30(14):3059–66.

3. Li W, Godzik A. Cd-hit: a fast program for clustering and comparing large sets of protein or nucleotide sequences. Bioinformatics. 2006 Jul 1;22(13):1658–9.

4. Minh BQ, Schmidt HA, Chernomor O, Schrempf D, Woodhams MD, von Haeseler A, et al. IQ-TREE 2: New Models and Efficient Methods for Phylogenetic Inference in the Genomic Era. Mol Biol Evol. 2020 May 1;37(5):1530–4.

5. Yu G. Using ggtree to Visualize Data on Tree-Like Structures. Curr Protoc Bioinformatics. 2020 Mar;69(1):e96.

6. Takada K, Kawakami C, Fan S, Chiba S, Zhong G, Gu C, et al. A humanized MDCK cell line for the efficient isolation and propagation of human influenza viruses. Nat Microbiol. 2019 Aug;4(8):1268–73.

7. Hoffmann E, Neumann G, Kawaoka Y, Hobom G, Webster RG. A DNA transfection system for generation of influenza A virus from eight plasmids. Proc Natl Acad Sci U S A. 2000 May 23;97(11):6108–13.

8. Neumann G, Watanabe T, Ito H, Watanabe S, Goto H, Gao P, et al. Generation of influenza A viruses entirely from cloned cDNAs. Proc Natl Acad Sci U S A. 1999 Aug 3;96(16):9345–50.

9. Liang W, Tan TJC, Wang Y, Lv H, Sun Y, Bruzzone R, et al. Egg-adaptive mutations of human influenza H3N2 virus are contingent on natural evolution. PLoS Pathog. 2022 Sep;18(9):e1010875.

10. Chan MC, Chan RW, Chan LL, Mok CK, Hui KP, Fong JH, et al. Tropism and innate host responses of a novel avian influenza A H7N9 virus: an analysis of ex-vivo and in-vitro cultures of the human respiratory tract. Lancet Respir Med. 2013 Sep;1(7):534–42.

11. Chan MC, Chan RW, Yu WC, Ho CC, Yuen KM, Fong JH, et al. Tropism and innate host responses of the 2009 pandemic H1N1 influenza virus in ex vivo and in vitro cultures of human conjunctiva and respiratory tract. Am J Pathol. 2010 Apr;176(4):1828–40.

12. Organization WH. SEROLOGICAL DIAGNOSIS OF INFLUENZA BY MICRONEUTRALIZATION ASSAY. [cited; Available from: https://www.who.int/publications-detail-redirect/serological-diagnosis-of-influenza-by-microneutralization-assay

